# Better data, better trees: GenBank-GISAID deduplication and source-specific artifact masking in viral genomics

**DOI:** 10.64898/2026.06.12.731931

**Authors:** Laise de Moraes, Andrêza Leite de Alencar, Marius Brusselmans, Darlan da S. Candido, Nuno R. Faria, Simon Dellicour, Philippe Lemey, Ricardo Khouri

## Abstract

GenBank and GISAID are the primary repositories for viral genomic data, but integrating records across them remains a challenge. The same sequence could be made available in both databases without any cross-reference linking the two entries. Consequently, there is no systematic way to identify this redundancy, which compromises the compilation of representative, non-redundant large-scale datasets. In parallel, the growth of viral genomic data has increased the risk of systematic technical artifacts introduced during sequencing or assembly. These artifacts can inflate substitution rate estimates and degrade temporal signal, biasing evolutionary rate estimates. To address both challenges, here we present a formal, reproducible workflow integrating two newly developed complementary tools: G2G matcher for cross-repository harmonization and Lab-Specific Bias FILTer (LSBFILT) for masking of laboratory-specific artifacts. Using the Eastern/Central/South African (ECSA) chikungunya virus lineage as a proof-of-concept, we demonstrate that our integrated workflow restores temporal signal and provides a robust, curated dataset for downstream phylodynamic analyses. Critically, restricting masking of homoplastic sites to specific sequences reduces the substitution rate estimate from an inflated 8.517 × 10^−4^ to 5.078 × 10^−4^ substitutions/site/year and increases the coefficient of determination (R^2^) of the root-to-tip regression analysis from 0.353 to 0.677. By enabling systematic cross-repository harmonization and source-specific artifact masking, we provide the molecular epidemiological community with scalable tools to reconcile fragmented genomic data and reduce technical biases, fostering more accurate and reproducible phylogenetic analysis. G2G matcher is available at https://github.com/andrezaleite/G2G-Matcher, and LSBFILT at https://github.com/khourious/LSBFILT.

## Introduction

Genomic sequencing is a powerful tool for molecular epidemiology of viral infections, allowing us to identify new viruses (Wu et al. 2020; Zhou et al. 2020), characterize emerging lineages and recombinant forms (Greninger et al. 2010; Faria et al. 2021; Naveca et al. 2024), trace their origins and natural reservoirs (Grubaugh et al. 2018; Pekar et al. 2025), reconstruct their dispersal history and dynamics across spatial and temporal scales (Dudas et al. 2017; Faria et al. 2017; Kraemer et al. 2021; De Moraes et al. 2023), and to test hypotheses about the external factors and intervention strategies impacting their spread (Lemey et al. 2021; Dellicour et al. 2025); with investigations ranging from localized outbreaks (Weaver 2014) to broader settings (Worobey et al. 2016; Santa Ardisson et al. 2025).

Viral genomic data are distributed across multiple databases, with GenBank (Benson et al. 2014), GISAID (Khare et al. 2021) and, more recently, Pathoplexus (Dalla Vecchia 2024) representing the most prominent resources in molecular epidemiological research. Pathoplexus functions as an upstream data-sharing platform, with sequences ultimately submitted to GenBank. However, GenBank and GISAID operate under distinct data-sharing frameworks and metadata standards, leading to partial and asymmetric overlap between repositories. Consequently, the same viral isolate is frequently represented across both databases with non-identical identifiers and attribute fields while each repository retains unique records and incomplete cross-referencing limits straightforward integration.

Large genomic datasets generated during outbreaks, often assembled from multiple sequencing efforts, are particularly susceptible to batch effects arising from laboratory-specific biases. These include variation in library preparation methods (*e*.*g*., amplicon-based approaches employing distinct primer schemes, probe-capture enrichment, or shotgun metagenomics), sequencing technologies (*e*.*g*., short-read and long-read platforms), and genome assembly workflows (*e*.*g*., choice of assembly algorithm, reference-guided mapping, or *de novo* assembly). Such technical heterogeneity can systematically bias phylogenetic clustering patterns, inflate substitution rate estimates, and degrade temporal signal in phylodynamic analyses.

In the context of SARS-CoV-2, the massive global sequencing effort underscored the practical impact of these biases. During the SARS-CoV-2 Delta wave, dropout of an ARTIC V3 primer led to widespread artefactual variation at Spike position 142, which was initially misinterpreted as a recurrent adaptive mutation (Sanderson and Barrett 2021). Subsequent work has shown that laboratory-associated homoplastic sites can distort evolutionary reconstructions, while recurrent sequencing errors combined with mutation rate variation introduce widespread homoplasies even in pandemic-scale datasets comprising millions of genomes (Turakhia et al. 2020; De Maio et al. 2026; Hunt et al. 2026). Similar concerns have been raised in a study of picornaviruses, where error rates as low as ∼0.5% and/or mis-specified sampling dates shifted by only five years were shown to substantially destabilize phylogenetic inference and increase the variance of evolutionary parameter estimates (Vakulenko et al. 2019).

As part of an ongoing genomic surveillance effort, we conducted preliminary phylogenetic inference analyses of publicly available chikungunya virus (CHIKV; *Togaviridae*: *Alphavirus chikungunya*) genomes, and we observed that the Eastern/Central/South African (ECSA) CHIKV lineage exhibited an unexpectedly elevated substitution rate and compromised temporal signal, consistent with the presence of widespread technical artifacts. This made it an ideal model to investigate these issues.

Here, we present a formal, reproducible workflow that integrates two complementary tools: G2G matcher, for systematic de-duplication and harmonization of viral genomic metadata across GenBank and GISAID; and Lab-Specific Bias FILTer (LSBFILT), a pipeline for identifying and masking laboratory-derived, source-specific technical artifacts.

Using the CHIKV-ECSA lineage as a proof-of-concept, we demonstrate how the application of this integrated workflow restores temporal signal, reduces substitution rate inflation, and provides a robust, curated dataset for downstream phylodynamic analyses.

### Building a Cross-Database Integration Strategy to Resolve Redundancy in GenBank and GISAID Viral Genomic Metadata

We developed G2G matcher, a systematic multi-layered framework designed to enable the consolidation of genomic metadata across GenBank (Benson et al. 2014) and GISAID (Khare et al. 2021). The tool executes a structured workflow comprising data harmonization, multi-strategy matching, and filtering to provide a non-redundant dataset optimized for downstream phylodynamic and molecular evolution studies. G2G matcher was implemented in Python (Rossum and Drake 2010), utilizing pandas (McKinney 2010) for data manipulation, Biopython (Cock et al. 2009) for sequence handling, and tqdm (Casper da Costa-Luis et al. 2026) for progress monitoring. The complete framework is provided as a Jupyter Notebook (Kluyver Thomas et al. 2016) and available at https://github.com/andrezaleite/G2G-Matcher.

Initial processing involves filtering sequences based on a minimum length threshold; reconciling location names and parsing administrative hierarchies into Country, State/Region, and City/Municipality levels; standardizing collection date to the ISO 8601 format (YYYY-MM-DD) with support for partial date entries; and extracting isolate identifiers from GenBank record descriptions (definition fields) using customized Regular Expression (Regex) patterns.

Following this preprocessing, the core integration logic utilizes three complementary strategies to identify corresponding records between repositories, and assigns confidence scores based on the degree and type of supporting evidence: (a) FASTA-based matching, which performs direct comparison of nucleotide sequences; (b) isolate field-based fuzzy matching, which leverages a semantic matching approach via the RapidFuzz library (Ye et al. 2021) to compare isolate names and extracted identifiers, with a 80% similarity threshold that accounts for minor orthographic variations or naming conventions across platforms; and (c) metadata-based matching, an exact matching approach focusing on the intersection of location, collection date, and sequence length. FASTA-based matches receive priority, followed by isolate field-based matches, and metadata-based matches.

To eliminate conflicting assignments, G2G matcher employs a bidirectional de-duplication approach, in which primary matching proceeds from GenBank to GISAID, followed by reciprocal GISAID to GenBank resolution. Each record is assigned an integration status:

- MATCH: High-confidence identity supported by nucleotide sequence data and/or multiple metadata overlaps.
- HIGH_POTENTIAL_MATCH: High evidence (typically nucleotide sequence identity) requiring minor validation.
- POTENTIAL_MATCH: Metadata overlap without sequence identity, or lower-score fuzzy matches.
- GENBANK_ONLY / GISAID_ONLY: Unique records present in only one database.

To ensure biological relevance, the G2G matcher automatically excludes experimental sequences, including synthetic constructs, patent-related sequences, vaccine-derived strains, animal models, and laboratory-adapted lineages. The structured output provides parallel columns distinguished by repository-specific prefixes *‘GenBank_’* and *‘GISAID_’*, alongside integration status (*‘MATCH’, ‘HIGH_POTENTIAL_MATCH’, ‘POTENTIAL_MATCH’, ‘GENBANK_ONLY’, ‘GISAID_ONLY’*) and supporting evidence (*‘FASTA’, ‘ISOLATE’, ‘LCDL’*).

We emphasize that the G2G consolidated dataset provides a reference point for manual curation, an essential step that this pipeline does not replace. Manual review remains critical for validating ambiguous matches, verifying metadata accuracy, and resolving conflicts that cannot be resolved algorithmically, such as distinguishing between genuine duplication and independently sampled isolates from the same spatiotemporal context.

### Compiling a Cross-Repository Dataset

We compiled a CHIKV dataset comprising near-full genome (>7,000 bp) sequences retrieved from GenBank and GISAID on March 29, 2026. The integration process allowed cross-repository verification of methodological details, including library preparation approaches (*e*.*g*., amplicon with their associated primer sets, shotgun, hybrid capture), sequencing technologies, and sequencing laboratory and their respective countries. Information about the submitting laboratory was retrieved from GISAID based on the EPI_SET Identifier using the GEDEX utility (https://github.com/khourious/GEDEX).

We then filtered for the ECSA lineage using the Chikungunya Virus Typing Tool v3.72 implemented in Genome Detective Virus Tool v2.104 (Fonseca et al. 2019; Vilsker et al. 2019) to generate a CHIKV-ECSA dataset. We subsequently filtered genomes to exclude sequences with missing collection dates and/or sampling locations that could not be confirmed, along with sequences that could not be manually validated by cross-referencing between GenBank and GISAID records, resulting in a final CHIKV-ECSA dataset of 8,617 genomes. Figure 1 provides a detailed flowchart illustrating the key steps of the CHIKV-ECSA dataset compilation.

**Figure 1.**
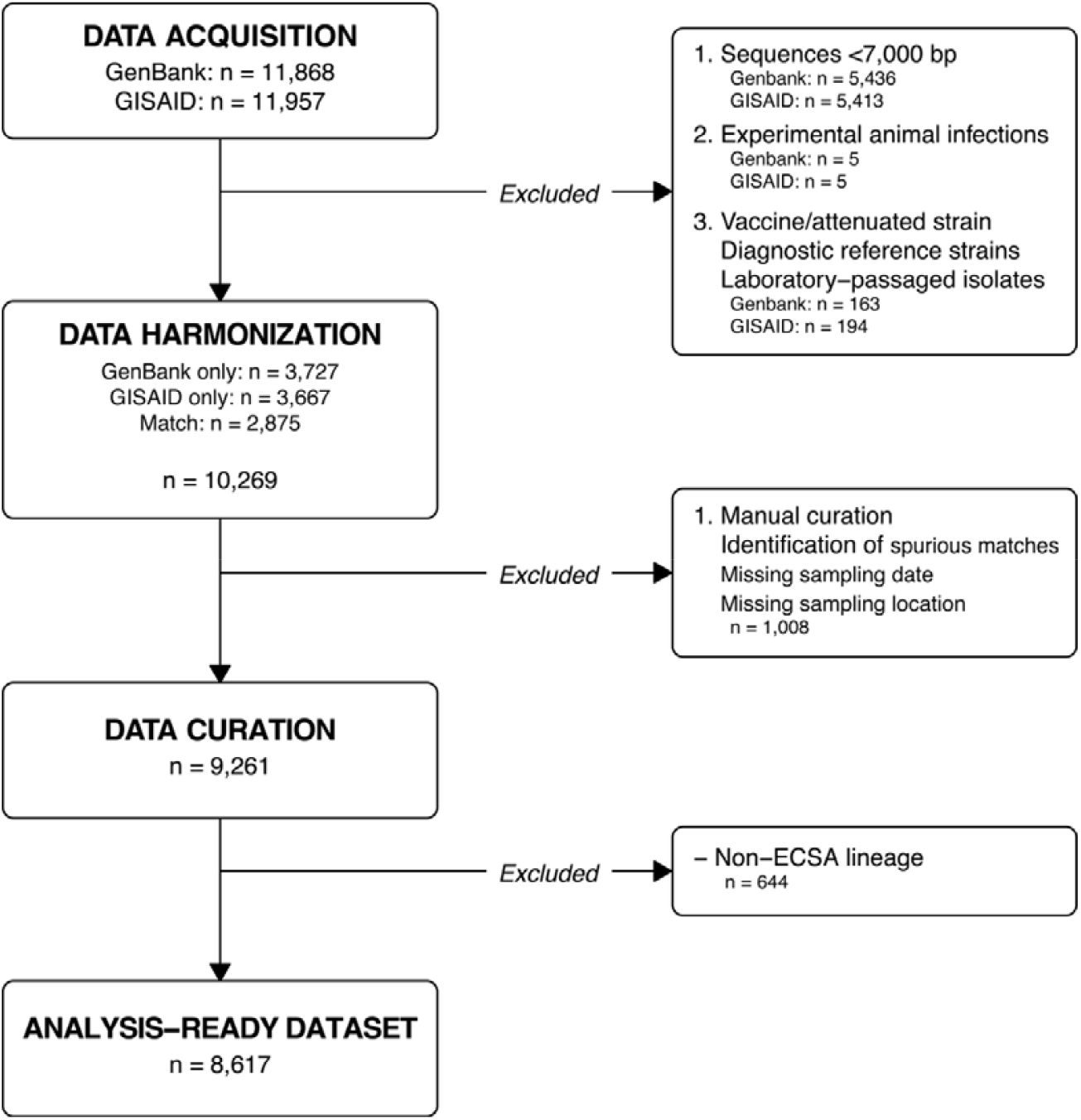
Flowchart of CHIKV-ECSA dataset compilation. Data acquisition comprised 11,868 GenBank and 11,957 GISAID records. Data harmonization yielded 10,269 sequences after merging both repositories and excluding those <7,000 bp and/or from non-natural sources. Data curation retained 9,261 sequences after removed spurious matches and/or records with incomplete metadata record. Subsequent lineage assignment producing the final ‘analysis-ready’ dataset of 8,617 CHIKV-ECSA genomes. In total, 2,490 sequences were shared between databases, while 3,056 were unique to GenBank and 3,071 to GISAID.

We then performed a multiple alignment using MAFFT v7.526 (Katoh 2002; Katoh and Standley 2013) that included the reference NC_004162.2, followed by minimal manual curation using AliView v1.31 (Larsson 2014). We performed phylogenetic reconstruction using the maximum likelihood phylogenetic inference method implemented in FastTree v2.2.0 (Price et al. 2010), under the CAT-GTR+G20 model, with pairwise distances initially calculated using the Jukes-Cantor model with balanced joins for initial topology construction. We assessed node support using 1,000 SH-like replicates while performing two rounds of NNI and SPR moves during tree searches, alongside maximum-likelihood NNI optimization at each iteration.

### Implementation of the Laboratory-Specific Bias Filter Pipeline

To identify technical artifacts and laboratory-specific biases in the CHIKV-ECSA dataset, we implemented a modified version of the workflow originally developed by Turakhia and colleagues to identify problematic positions in SARS-CoV-2 genomes. Our analytical pipeline is available at https://github.com/khourious/LSBFILT.

Turakhia and colleagues (2020) established a heuristic approach that combines phylogenetic parsimony evaluation with metadata from sequencing laboratories. Briefly, their approach first employs a phylogenetic parsimony analysis to identify sites at which the same mutation arises independently multiple times on different branches of the evolutionary tree, indicating high homoplasy inconsistent with standard substitution dynamics over relatively short time scales. Subsequently, sites are cross-referenced with laboratory metadata, and alleles showing substantial bias associated with a single laboratory source are masked as probable systematic artifacts. They further examined linkage disequilibrium among alleles, which measures how frequently two mutations are inherited together on the same genome and evaluated overlap with primer binding sites to characterize other potential technical artifacts. Based on these criteria, Turakhia et al. (2020) recommended global masking of laboratory-associated and highly homoplastic positions (the same positions masked in all sequences) prior to phylogenetic inference to mitigate bias in downstream analyses.

Here, we extended this pipeline by allowing adjustments of parsimony, laboratory bias, and linkage disequilibrium filters. In addition, we implemented an allele level, source-specific masking workflow to enable artifact detection across diverse viral genomic datasets. The implementation requires four inputs: (i) a multiple sequence alignment in FASTA format containing both viral sequences and the reference genome; (ii) a phylogenetic tree in NEWICK format generated from the alignment; (iii) a TSV-formatted metadata table containing ‘*sequence_id*’, *‘sequencing_lab’, ‘sequencing_lab_country’*, and ‘*sequencing_lib_prep*’; and (iv) a directory containing tab-formatted text files with genomic coordinates (BED format) for primer schemes, each named to match the primer scheme specified in the *‘sequencing_lib_prep’* field. To support future applications, the BED files compiled for the CHIKV-ECSA dataset are available at https://github.com/khourious/LSBFILT/BED/CHIKV.

Initial processing involves the generation of a VCF file from the multiple alignment using snp-sites v2.5.1. Ambiguous base calls, including ‘N’ and other IUPAC ambiguity codes, are retained unchanged in the workflow (referred to as Unresolved). To assess how ambiguity-handling approaches influence artifact detection, ambiguous bases were either replaced with the reference allele (Ref Resolved) or assigned the most plausible alternative allele (Alt Resolved). Parsimony scores for all three approaches were computed using the ‘*find_parsimonious_assignments’* module of the strain phylogenetics (Turakhia et al. 2020).

Variable positions were subsequently selected based on a defined parsimony score threshold and the proportion of alternate allele calls linked to each sequencing laboratory and corresponding country was calculated. Variable positions were flagged as laboratory-associated when a given percentage of sequences carrying the alternate allele originated from a single source. The ratio of the parsimony score (PS) to alternate allele count (AC), defined as PS/AC, was calculated. Ratios ≥ 0.5 indicate variable positions with high recurrence across the phylogeny but low population frequency, a pattern suggestive of non-heritable and technically derived artifacts rather than true evolutionary events.

Allele positions were selected for masking if they exhibited a PS/AC ratio ≥ 0.5 in at least two of the three ambiguity-handling workflows (Unresolved, Ref Resolved, and Alt Resolved). In contrast to global masking strategies, this approach implements a conservative, source-specific masking workflow, restricting masking to sequences identified as laboratory-or laboratory-country-associated. By removing artifacts only where they are most likely to occur, this masking approach strikes a balance between minimizing the misinterpretation of technical artifacts as evolutionary signal in phylogenetic reconstructions and preserving authentic biological variation elsewhere in the alignment.

Variable positions were also annotated as primer-associated if they overlapped or fell within the vicinity of primer binding sites (±10 bp), using BED files corresponding to the library preparation method applied to each sequence. Sequences lacking information on the library preparation method or generated by methods other than amplicon sequencing (*e*.*g*., shotgun or hybrid capture) were excluded from this analysis, underscoring the importance of complete methodological metadata for artifact identification. To assess linkage disequilibrium between variable positions, we calculated coefficient of determination (R^2^) values using PLINK v2.0.0a.6.9 (Purcell et al. 2007), retaining those meeting a predefined R^2^ threshold as potentially linked technical artifacts.

Finally, tables were generated detailing parsimony scores, allele frequencies, laboratory associations, primer overlaps, linkage disequilibrium patterns, and masking decisions. These outputs provide full transparency and allow researchers to review, validate, or adjust the filtered variable positions as needed for downstream analyses.

### Assessing the Impact of Lab-Specific Masking on Phylogenetic Signal and Topological Stability

To identify the most appropriate masking strategy for a given dataset, we evaluated the effect of masking laboratory-or laboratory country-associated variable positions on temporal signal and topological stability. We performed phylogenetic reconstruction from the unmasked alignment (referred to as the baseline) and from alignments masked under different combinations of minimum parsimony scores (≥ 1-10) and minimum specific bias association (≥ 60-90%) thresholds, applied for two independent criteria: sequencing laboratory (*e*.*g*., PS1<u>LSB</u>10) and sequencing laboratory country (*e*.*g*., PS1<u>LCSB</u>10). All trees were reconstructed using FastTree as previously described. For each masking strategy, we evaluate the temporal signal by performing linear regression of root-to-tip (RTT) distance against sampling dates, using the root that maximized the R^2^ of the linear regression.

We then select the top 3 masking strategies that simultaneously maximize R^2^, mask a total of less than 1,000 alleles (to preserve biological signal), and reduce the substitution rate (since technical artifacts artificially inflate rate estimates). To assess their effect on tree topology, we generated nine additional independent phylogenetic replicates, reconstructed using FastTree as previously described. Two distance metrics were computed to capture complementary aspects of topological divergence: Robinson-Foulds (RF) (Robinson and Foulds 1981), which evaluates whether the same sets of sequences cluster together, without considering branch lengths; and Path Difference (PD) (Steel and Penny 1993), which measures how much the distances between all pairs of sequences differ when branch lengths are taken into account.

To select the single best among the top three masking strategies, we removed outliers with studentized residuals > 3 and repeated the RTT regression. The strategy yielding the highest post-removal R^2^ was chosen as the final dataset. To characterize topological differences between Baseline and the selected masking strategy, we generated a tanglegram. For legibility, the dataset was subsampled to 20% by retaining every 5th tip as ordered in the Baseline tree topology following branch sorting, and the same subset of sequences was then applied to the selected masking strategy tree. Trees were untangled over ten iterative rounds to minimize crossing lines between neighbors. Tree parsing, manipulation, and visualization were performed using baltic v0.3.0 (Dudas 2024), together with matplotlib v3.10.9 (Hunter 2007), pandas v2.2.2 (McKinney 2010), and numpy v2.4.4 (Harris et al. 2020) in a Jupyter Notebook v4.5.6 (Kluyver Thomas et al. 2016).

All remaining statistical analyses, phylogenetic assessments, and visualizations described in this section were performed using R Statistical Software v4.5.2 (R Core Team 2025) within the RStudio environment v2026.01.1+403 (Posit Team 2026). We performed RTT regression and assessed topological relationships using “ape” v5.8.1 (Paradis and Schliep 2019), “MASS” v7.3.65 (Venables and Ripley 2002), “phangorn” v2.12.1 (Schliep 2011; Schliep et al. 2017), “phytools” v2.5.2 (Revell 2024), and “seqinr” v4.2.36 (Bastolla 2007). Data processing was assessed with “dplyr” v1.2.0 (Wickham, François, et al. 2014), “glue” v1.8.0 (Hester and Bryan 2017), “lubridate” v1.9.5 (Grolemund and Wickham 2011), “readr” v2.2.0 (Wickham et al. 2015), “readxl” v1.4.5 (Wickham and Bryan 2015), “stringr” v1.6.0 (Wickham 2009), and “tidyr” v1.3.2 (Wickham, Vaughan, et al. 2014). Visualizations were generated using “cowplot” v1.2.0 (Wilke 2015), “ggnewscale” v0.5.2 (Campitelli 2019), “ggplot2” v4.0.2 (Wickham 2016), “ggrepel” v0.9.7 (Slowikowski 2016), “Gmisc” v3.2.0 (Gordon 2015), “grid” v4.5.2 (R Core Team 2025), “gridExtra” v2.3 (Auguie 2010), “scales” v1.4.0 (Wickham et al. 2011), and “scatterplot3d” v0.3.45 (Ligges and Mächler 2003).

### Identifying the Optimal Masking Strategy for the CHIKV-ECSA Dataset

We first assessed the temporal signal of the unmasked alignment. The baseline tree yielded an R^2^ value of 0.353 and a substitution rate of 8.517 × 10^−4^ substitutions/site/year (s/s/y; Figure 2). This R^2^ value indicates that only 35% of the variation in genetic distances is explained by sampling time, leaving the majority of the variance driven by other factors (Rambaut et al. 2016). The observed substitution rate is approximately two to three times higher than previously published estimates for CHIKV-ECSA, with mean substitution rates ranging from 2.30 × 10^−4^ to 4.40 × 10^−4^ s/s/y and 95% highest posterior density (HPD) intervals spanning 1.37 × 10^−4^ to 4.80 × 10^−4^ (Volk et al. 2010; Nunes et al. 2015; Souza et al. 2019; Deeba et al. 2020; Krambrich et al. 2024).

**Figure 2.**
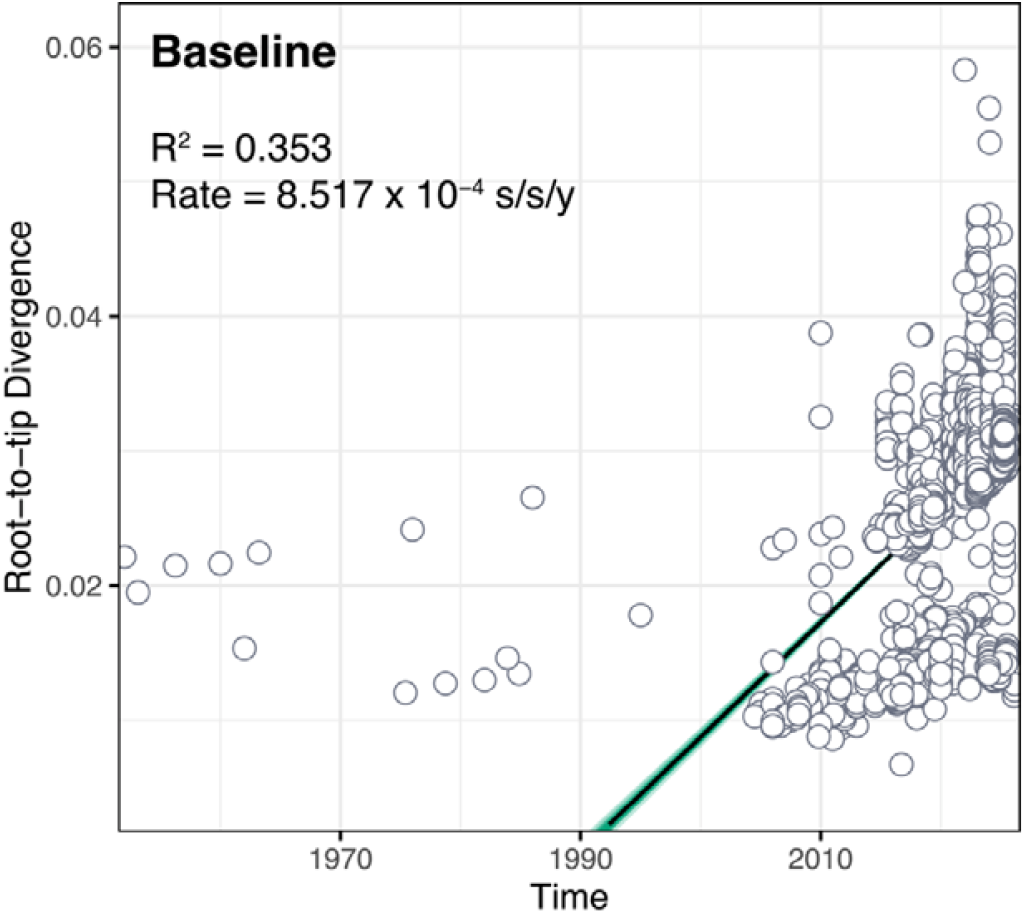
Root-to-tip regression of the baseline. The scatter plot shows root-to-tip genetic divergence as a function of sampling time, with each point representing a sequence, the black line showing the linear regression, and the green lines indicating the associated confidence interval.

Visual inspection of the RTT regression plot confirmed this interpretation, revealing a bimodal distribution with a large cluster of sequences deviating substantially from the main temporal trend, suggestive of a distinct artifact-afflicted subpopulation rather than isolated outliers (Figure 2).

A straightforward approach to mitigate these artifacts would be to exclude problematic sequences. Outlier exclusion is a standard practice in phylogenetic dataset construction, commonly applied to mitigate the influence of sequences with anomalous evolutionary signals that can distort temporal signal estimation and branch length inference (Rambaut et al. 2016; Lozano-Fernandez 2022). However, large-scale sequence exclusion risks losing valuable information about key introductions and critical events in regional transmission dynamics (Bar-Hen et al. 2008; Karcher et al. 2016; Villabona-Arenas et al. 2020).

To balance this trade-off and mitigate the inflation of temporal signal, we first applied source-specific masking before any sequence exclusion. We evaluated 80 masking strategy combinations, spanning ten minimum parsimony scores (1 to 10) and four minimum specific bias association thresholds (60%, 70%, 80%, and 90%), applied separately to laboratory-or laboratory-country-association (Supplementary Table 1). From these, the top 3 masking strategies according to our selection criteria (R^2^ maximization, ≤ 1,000 masked alleles, and substitution rate minimization) were: PS8LCSB60, corresponding to minimum parsimony score ≥ 8 and laboratory-country-specific bias ≥ 60%; PS8LCSB70 (≥ 8, ≥ 70%); and PS10LCSB60 (≥ 10, ≥ 60%) (Figure 3).

**Table 1.** Root-to-tip regression metrics for the Baseline and top three masking strategies applied to the CHIKV-ECSA dataset.

| Masking strategy | Before outlier exclusion |  |  | After outlier exclusion |  |  |
| --- | --- | --- | --- | --- | --- | --- |
| | Masked alleles (*n*) | Substitution rate (s/s/y) | $R^2$ | Removed sequences (*n*, %) | Substitution rate (s/s/y) | $R^2$ |
| Baseline | 0 | $8.517 \times 10^{-4}$ | 0.353 | - | - | - |
| PS8LCSB60 | 521 | $5.026 \times 10^{-4}$ | 0.522 | 136 (1.58) | $5.036 \times 10^{-4}$ | 0.669 |
| PS8LCSB70 | 496 | $4.844 \times 10^{-4}$ | 0.524 | 147 (1.71) | $4.835 \times 10^{-4}$ | 0.663 |
| **PS10LCSB60** | **379** | **$5.043 \times 10^{-4}$** | **0.543** | **145 (1.68)** | **$5.078 \times 10^{-4}$** | **0.677** |

Metrics are shown for the Baseline and top three masking strategies before and after the exclusion of outlier sequences (studentized residual > 3). Masking strategy nomenclature follows the pattern PS[minimum parsimony score][source type][minimum bias %]. All datasets prior to masking comprised 8,617 CHIKV-ECSA genomes. Masked alleles refer to positions across the reference NC\_004162.2 (11,826 nt) that were replaced with N in source-associated sequences. The optimal masking strategy selected is highlighted in bold. s/s/y: substitutions/site/year.

**Figure 3.**
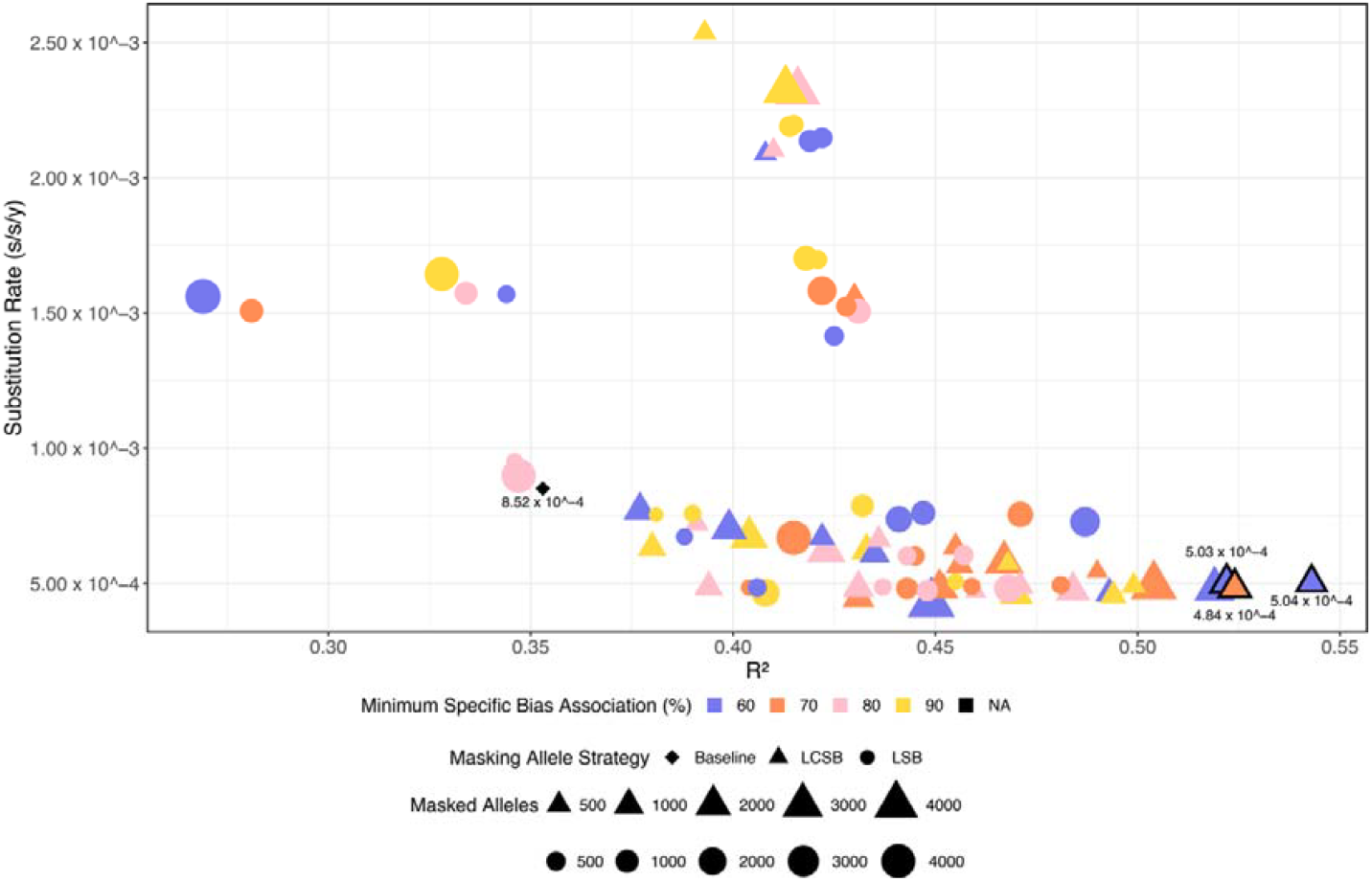
Effect of allele masking strategy on root-to-tip regression in the CHIKV-ECSA dataset. The scatter plot shows the relationship between root-to-tip temporal signal (coefficient of determination R^2^ from the root-to-tip regression) and substitution rate across 80 masking strategies applied to the CHIKV-ECSA dataset. Point shape indicates the masking criterion: LSB (laboratory-specific bias, circles), LCSB (laboratory-country-specific bias, triangles), and Baseline (unmasked alignment, diamond). Point colour denotes the minimum specific bias association threshold (blue, 60%; orange, 70%; pink, 80%; yellow, 90%). Point size reflects the number of masked alleles. The top three masking strategies selected for downstream analysis are outlined in black.

All three strategies substantially outperformed the Baseline, with R^2^ values ranging from 0.522 to 0.543 and substitution rates from 4.844 × 10^−4^ to 5.043 × 10^−4^ s/s/y (Table 1, Figure 4). Between 379 and 521 alleles were masked per strategy. The criteria used to mask each allele are detailed in Supplementary Tables 2 to 4. Overall, this indicates that some mutations are overrepresented in sequences from a single lab or country, and that these mutations are likely technical artefacts.

**Figure 4.**
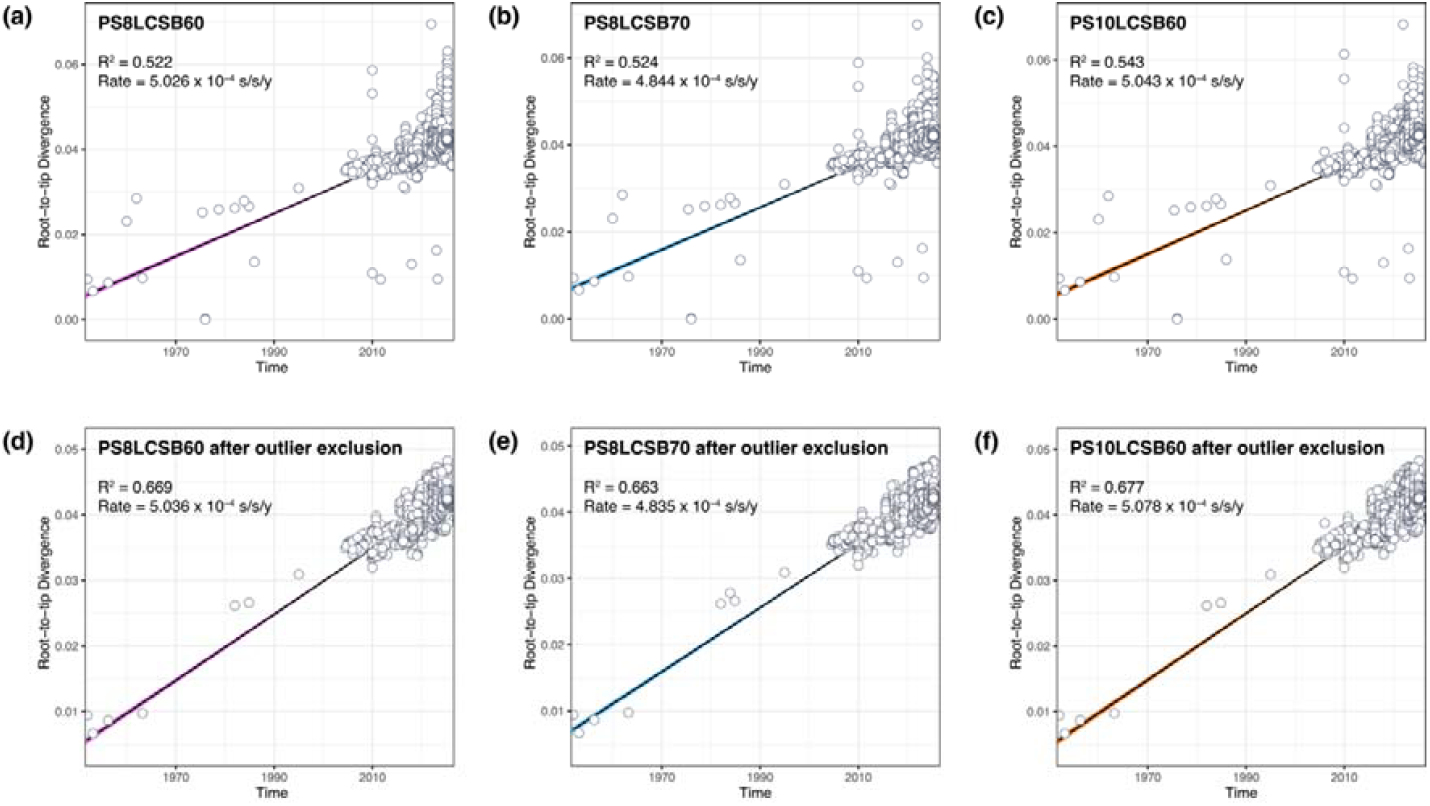
Root-to-tip regressions for the top three masking strategies before and after outlier exclusion. The scatter plot shows root-to-tip genetic divergence as a function of sampling time for PS8LCSB60 (A, before outlier removal; D, after), PS8LCSB70 (B, before; E, after), and PS10LCSB60 (C, before; F, after), with each point representing a sequence, the black line showing the linear regression, and colored lines indicating the confidence interval. Outliers with studentized residuals > 3 were removed prior to regression.

Regarding the effect on tree topology, multidimensional scaling (MDS) projections of inter-tree RF and PD distances recovered four well-defined, non-overlapping clusters in tree space, indicating highly reproducible topologies, with the masked conditions clustering more closely together than to the baseline, consistent with the shared effect of removing source-specific homoplastic signal (Figure 5).

**Figure 5.**
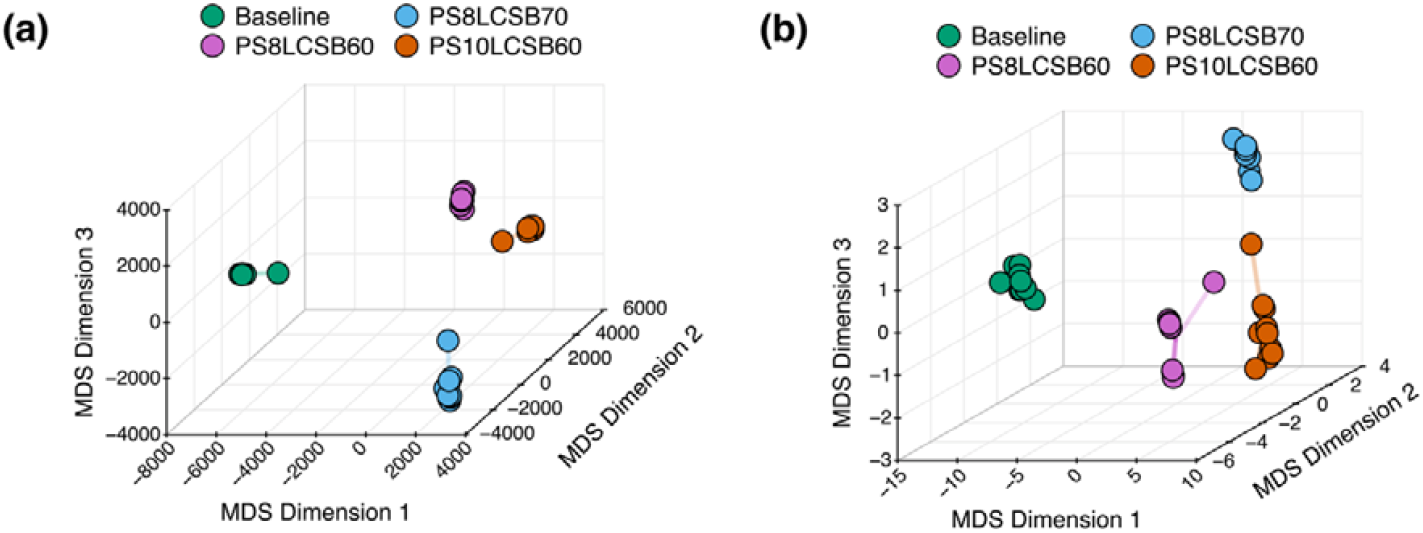
Topological distances between the Baseline and the top three masking strategies. Topological divergence was assessed by multidimensional scaling (MDS) of (A) Robinson-Foulds (RF) and (B) Path Difference (PD) across ten independent phylogenetic replicates. Points represent individual tree replicates in MDS space, with Baseline in green, PS8LCSB60 in pink, PS8LCSB70 in blue, and PS10LCSB60 in orange. Proximity in MDS space reflects topological similarity, and clustering indicates reproducible topologies.

The three masking strategies produced comparably strong improvements in temporal signal, suggesting that any of them would likely yield reliable results regardless of the final choice. To select a single representative, we used outlier removal as a final decision criterion. Unlike the large-scale sequence exclusion discussed earlier, we removed the remaining outliers, since the masking strategy has already resolved most artifacts. After removing outliers (studentized residuals > 3), the three strategies led to the exclusion of 136-147 sequences, with R^2^ values ranging from 0.663 to 0.677 and substitution rates from 4.835 × 10^−4^ to 5.078 × 10^−4^ s/s/y (Table 1 and Figure 4). We therefore selected PS10LCSB60 as the optimal masking strategy in our case, as it achieved the highest post-removal R^2^ (0.677).

Tanglegram comparison showed that topological disagreement between the baseline and PS10LCSB60 was concentrated within specific clades, most notably those comprising sequences from Brazil and La Réunion, the two countries whose sequencing efforts contributed the largest number of sequences to this dataset (Brazil: 48.91%, 4,215/8,617; La Réunion: 36.22%, 3,121/8,617). Sequences generated by the USA, Paraguay, China, and France were also affected, though only when they branched together with those from Brazil or La Réunion. This pattern was particularly evident across the 2020-2025 genomes (Figure 6).

**Figure 6.**
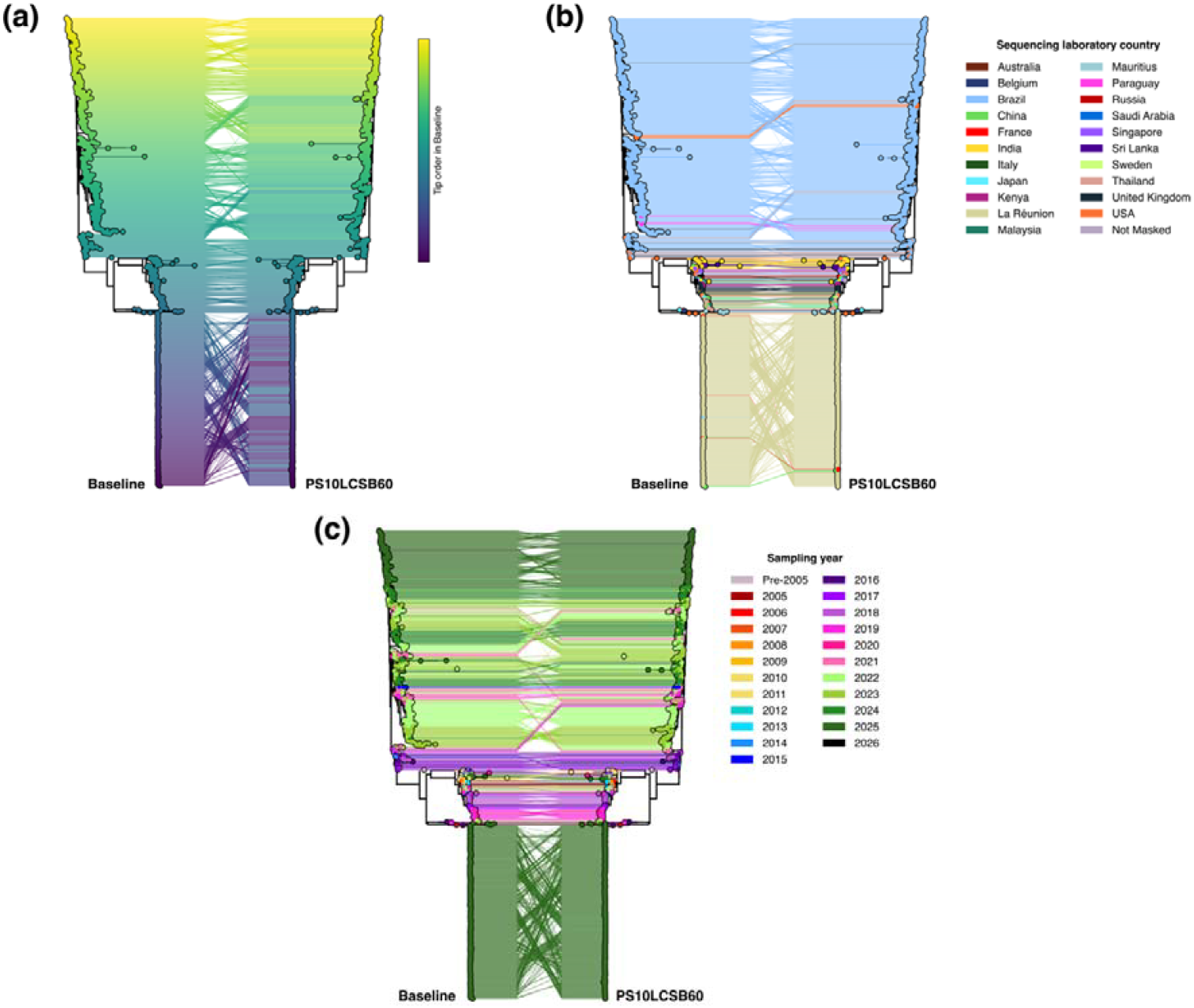
Topological comparison between Baseline and PS10LCSB60. The tanglegram shows topological relationships between the Baseline (left tree) and PS10LCSB60 (right tree), with connecting lines linking corresponding tips; line crossings indicate topological disagreement between the two trees. Tips are colored to illustrate different features, with (A) showing tip order in the Baseline tree as a gradient from first (yellow) to last (purple), to visualize tip repositioning in the PS10LCSB60 masking strategy; (B) showing sequencing laboratory country; and (C) showing sampling year. For legibility, trees were subsampled to 20% of tips and untangled over ten iterative rounds to minimize crossing lines between neighbors.

## Conclusion and Future Directions

The exponential growth of viral genomic data across multiple repositories provides an unprecedented empirical foundation for molecular epidemiological research while also increasing the risk of systematic technical artifacts that can distort phylogenetic inference. Our results provide compelling evidence for the existence of lab- and country-associated artefacts, support the view that heterogeneity in sequencing and analysis approaches is a key contributing factor, and propose a practical and scalable strategy to address these biases. We present two computational resources to address these challenges.

First, the G2G matcher streamlines harmonization of genomic data from GenBank and GISAID, addressing a critical gap in viral genomics where cross-repository integration has remained a manual and error-prone process. Second, the LSBFILT pipeline enables researchers to identify and mitigate laboratory-or laboratory-country-specific artifacts.

Using CHIKV-ECSA as a proof-of-concept, we demonstrate that restricting the filtering of homoplastic sites only to sequences derived from the source laboratory or laboratory country resolves a pronounced inflation of RTT distances, reducing the substitution rate estimate from an inflated 8.517 × 10^−4^ to a biologically plausible 5.078 × 10^−4^ s/s/y, consistent with prior estimates of CHIKV substitution rates (Volk et al. 2010; Nunes et al. 2015; Souza et al. 2019; Deeba et al. 2020; Krambrich et al. 2024). The increasing the R^2^ value from 0.353 to 0.677 improves the reliability of molecular clock analyses, and indicates that sampling time now explains most of the variation in genetic distances (Rambaut et al. 2016; Rieux and Balloux 2016), rather than stochastic or systematic sources of noise (Vakulenko et al. 2019). The masking strategy improved molecular clock reliability across all three CHIKV-ECSA sublineages, with the most pronounced effect observed in ECSA-2, the sublineage that dominates recent epidemic waves in the Americas and contributes with the largest number of sequences to the CHIKV-ECSA dataset (detailed results provides in the Supplementary Material). Our approach offers a middle ground between large-scale sequence removal, which can discard valuable outbreak data, and global site masking, which removes biological signal from all sequences.

Based on these analyses, we propose an enhanced phylogenetic workflow that combines established best practices and commonly used tools with the inclusion of LSBFILT for source-specific artifact masking. As a practical reference, and overview of this workflow is provided in Supplementary Table 5.

Our findings also underscore the importance of standardized deposition of raw sequencing reads and comprehensive metadata, which at present are often optional or inconsistently reported across public repositories. We advocate for the deposition of raw sequence reads (FASTQ, and where applicable, FAST5 or POD5) in public repositories such as the NCBI Sequence Read Archive (SRA) or the European Nucleotide Archive (ENA) as a fundamental requirement for ensuring reproducibility, transparency, and quality in genomic research (Brown et al. 2018; Byrd et al. 2020), particularly in epidemiological surveillance and public health contexts (Black et al. 2020).

The availability of raw data enables the scientific community to verify whether a reported variant represents a genuine biological mutation or a technical artifact, as well as to re-analyze data with new tools or improved parameters, facilitating discoveries unforeseen by the original data generators (Salfati et al. 2019; Goig et al. 2020). For instance, a comparative analysis of eight *de novo* assemblers on ^∼^400 SARS-CoV-2 samples reported that at least 9% of identified variants were assembler-specific (Islam et al. 2021).

Equally important is the standardized reporting of metadata, including sampling strategy, library preparation method, sequencing platform, coverage depth, and both sequencing and assembly QC metrics. While this is more affordable than the massive deposition of raw genomic data, ensuring its proper implementation remains a significant challenge. In the context of the COVID-19 pandemic, Schriml et al. (Schriml et al. 2020) warned that the lack of metadata standardization reduces the near- and long-term utility of genomic data, underscoring that a clear description of the WHO, WHAT, HOW, WHERE, and WHEN is essential for comparative analysis, public health response, and outbreak progression assessment. Illustrating this point, an investigation of SARS-CoV-2 genomic metadata in the GISAID database found that ^∼^10% of “originating lab” and “submitting lab” entries contained spelling errors or inconsistent naming conventions. Such inconsistencies can cause mutations from a single laboratory to appear linked to multiple sources, masking laboratory-specific artifacts and undermining phylogenetic inference (Gozashti and Corbett-Detig 2021).

Together with raw data access, these would allow robust independent validation and strengthen the reliability of genomic analyses. Recent initiatives such as Pathoplexus (Dalla Vecchia 2024) have begun to address these gaps by implementing standardized metadata requirements and encouraging raw read deposition. However, Pathoplexus currently includes only a limited set of viral pathogens, and these practices remain the exception rather than the norm across the broader genomic data landscape.

As genomic surveillance expands globally and sequencing efforts become increasingly decentralized, the ability to mitigate source-specific biases becomes essential for accurate phylodynamic inference. The expansion of sequencing efforts thus not only improves outbreak detection and enables real-time monitoring of pathogen spread but also creates the statistical conditions necessary to detect and filter out systematic and stochastic errors that would otherwise remain hidden in homogeneous, centrally generated datasets. Given these considerations, we encourage future studies to extend detection to other batch effects (*e*.*g*., sequencing platforms, assembly approaches, library preparation kits, or linkage disequilibrium patterns) and to validate this approach across a broader range of viral pathogens with distinct evolutionary dynamics.

## Supplementary Material

Supplementary Material is available at *Molecular Biology and Evolution* online.

## Acknowledgments

We acknowledge all laboratories worldwide for the sequence data deposited in GenBank and GISAID EpiArbo. A full GISAID acknowledgement table can be found with the EPI_SET ID EPI_SET_260423vy (https://doi.org/10.55876/gis8.260423vy), in Data Acknowledgement Locator under GISAID Resources (https://www.gisaid.org). We gratefully acknowledge the researchers who shared detailed amplicon primer schemes upon request. The authors also thank the HPC Core Facility (RPT04A/FIOCRUZ) for computational resources. The manuscript was translated into English with the assistance of DeepSeek v3.2. The translated version was carefully reviewed to ensure conceptual fidelity and linguistic adequacy; the authors take full responsibility for the final content.

## Author Contributions

*Laise de Moraes:* Conceptualization, Data Curation, Formal Analysis, Software, Writing-Original Draft, and Writing-Review & Editing; *Andrêza Leite de Alencar:* Conceptualization, Software, and Writing-Review & Editing; *Marius Brusselmans:* Conceptualization, and Writing-Review & Editing; *Darlan S. Candido:* Writing-Review & Editing; *Nuno R. Faria:* Writing-Review & Editing; *Simon Dellicour:* Conceptualization, Resources, Writing-Review & Editing, and Supervision; *Philippe Lemey:* Conceptualization, Resources, Writing-Review & Editing, and Supervision; *Ricardo Khouri:* Conceptualization, Formal Analysis, Resources, Writing-Review & Editing, Project Administration, and Funding Acquisition. All authors have read and approved the manuscript.

## Data Availability

G2G matcher is available at https://github.com/andrezaleite/G2G-Matcher Lab-Specific Bias FILTer (LSBFILT) is available at https://github.com/khourious/LSBFILT. Supporting files for the bioinformatic analysis are available on Figshare at https://doi.org/10.6084/m9.figshare.32386650.

## References

Auguie B. 2010. gridExtra: Miscellaneous Functions for “Grid” Graphics. :2.3. Available from: https://CRAN.R-project.org/package=gridExtra

Bar-Hen A, Mariadassou M, Poursat M-A, Vandenkoornhuyse P. 2008. Influence Function for Robust Phylogenetic Reconstructions. Molecular Biology and Evolution 25:869–873.

Bastolla U ed. 2007. Structural approaches to sequence evolution: molecules, networks, populations. Berlinlll; New York: Springer

Benson DA, Clark K, Karsch-Mizrachi I, Lipman DJ, Ostell J, Sayers EW. 2014. GenBank. Nucl. Acids Res. 42:D32–D37.

Black A, MacCannell DR, Sibley TR, Bedford T. 2020. Ten recommendations for supporting open pathogen genomic analysis in public health. Nat Med 26:832–841.

Brown AV, Campbell JD, Assefa T, Grant D, Nelson RT, Weeks NT, Cannon SB. 2018. Ten quick tips for sharing open genomic data.Ouellette F, editor. PLoS Comput Biol 14:e1006472.

Byrd JB, Greene AC, Prasad DV, Jiang X, Greene CS. 2020. Responsible, practical genomic data sharing that accelerates research. Nat Rev Genet 21:615–629.

Campitelli E. 2019. ggnewscale: Multiple Fill and Colour Scales in “ggplot2.” :0.5.2. Available from: https://CRAN.R-project.org/package=ggnewscale

Casper da Costa-Luis, Stephen Karl Larroque, Kyle Altendorf, Hadrien Mary richardsheridan, Mikhail Korobov, Noam Raphael, Ivan Ivanov, Marcel Bargull, Nishant Rodrigues, et al. 2026. tqdm: A fast, Extensible Progress Bar for Python and CLI. Available from: https://zenodo.org/doi/10.5281/zenodo.595120

Cock PJA, Antao T, Chang JT, Chapman BA, Cox CJ, Dalke A, Friedberg I, Hamelryck T, Kauff F, Wilczynski B, et al. 2009. Biopython: freely available Python tools for computational molecular biology and bioinformatics. Bioinformatics 25:1422–1423.

Dalla Vecchia E. 2024. Pathoplexus: towards fair and transparent sequence sharing. The Lancet Microbe 5:100995.

De Maio N, Willemsen M, Martin S, Guo Z, Saha A, Hunt M, Ly-Trong N, Minh BQ, Iqbal Z, Goldman N. 2026. Rate variation and recurrent sequence errors in pandemic-scale phylogenetics. Nat Methods 23:565–573.

De Moraes L, Portilho MM, Vrancken B, Van Den Broeck F, Santos LA, Cucco M, Tauro LB, Kikuti M, Silva MMO, Campos GS, et al. 2023. Analyses of Early ZIKV Genomes Are Consistent with Viral Spread from Northeast Brazil to the Americas. Viruses 15:1236.

Deeba F, Haider MSH, Ahmed A, Tazeen A, Faizan MI, Salam N, Hussain T, Alamery SF, Parveen S. 2020. Global transmission and evolutionary dynamics of the Chikungunya virus. Epidemiol. Infect. 148:e63.

Dellicour S, Gámbaro F, Jacquot M, Lequime S, Baele G, Gilbert M, Pybus OG, Suchard MA, Lemey P. 2025. Comparative performance of viral landscape phylogeography approaches. Proc. Natl. Acad. Sci. U.S.A. 122:e2506743122.

Dudas G. 2024. baltic: Backronymed Adaptable Lightweight Tree Import Code. Available from: https://github.com/evogytis/baltic

Dudas G, Carvalho LM, Bedford T, Tatem AJ, Baele G, Faria NR, Park DJ, Ladner JT, Arias A, Asogun D, et al. 2017. Virus genomes reveal factors that spread and sustained the Ebola epidemic. Nature 544:309–315.

Faria NR, Mellan TA, Whittaker C, Claro IM, Candido DDS, Mishra S, Crispim MAE, Sales FCS, Hawryluk I, McCrone JT, et al. 2021. Genomics and epidemiology of the P.1 SARS-CoV-2 lineage in Manaus, Brazil. Science 372:815–821.

Faria NR, Quick J, Claro IM, Thézé J, De Jesus JG, Giovanetti M, Kraemer MUG, Hill SC, Black A, Da Costa AC, et al. 2017. Establishment and cryptic transmission of Zika virus in Brazil and the Americas. Nature 546:406–410.

Fonseca V, Libin PJK, Theys K, Faria NR, Nunes MRT, Restovic MI, Freire M, Giovanetti M, Cuypers L, Nowé A, et al. 2019. A computational method for the identification of Dengue, Zika and Chikungunya virus species and genotypes. Rodriguez-Barraquer I, editor. PLoS Negl Trop Dis 13:e0007231.

Goig GA, Blanco S, Garcia-Basteiro AL, Comas I. 2020. Contaminant DNA in bacterial sequencing experiments is a major source of false genetic variability. BMC Biol 18:24.

Gordon M. 2015. Gmisc: Descriptive Statistics, Transition Plots, and More. :3.2.0. Available from: https://CRAN.R-project.org/package=Gmisc

Gozashti L, Corbett-Detig R. 2021. Shortcomings of SARS-CoV-2 genomic metadata. BMC Res Notes 14:189.

Greninger AL, Chen EC, Sittler T, Scheinerman A, Roubinian N, Yu G, Kim E, Pillai DR, Guyard C, Mazzulli T, et al. 2010. A Metagenomic Analysis of Pandemic Influenza A (2009 H1N1) Infection in Patients from North America.Tripp R, editor. PLoS ONE 5:e13381.

Grolemund G, Wickham H. 2011. Dates and Times Made Easy with lubridate. J. Stat. Soft.

Grubaugh ND, Ladner JT, Lemey P, Pybus OG, Rambaut A, Holmes EC, Andersen KG. 2018. Tracking virus outbreaks in the twenty-first century. Nat Microbiol 4:10–19.

Harris CR, Millman KJ, Van Der Walt SJ, Gommers R, Virtanen P, Cournapeau D, Wieser E, Taylor J, Berg S, Smith NJ, et al. 2020. Array programming with NumPy. Nature 585:357–362.

Hester J, Bryan J. 2017. glue: Interpreted String Literals. :1.8.0. Available from: https://CRAN.R-project.org/package=glue

Hunt M, Hinrichs AS, Anderson D, Karim L, Dearlove BL, Knaggs J, Constantinides B, Fowler PW, Rodger G, Street T, et al. 2026. Addressing pandemic-wide systematic errors in the SARS-CoV-2 phylogeny. Nat Methods 23:653–662.

Hunter JD. 2007. Matplotlib: A 2D Graphics Environment. Comput. Sci. Eng. 9:90–95.

Islam R, Raju RS, Tasnim N, Shihab IH, Bhuiyan MA, Araf Y, Islam T. 2021. Choice of assemblers has a critical impact on de novo assembly of SARS-CoV-2 genome and characterizing variants. Briefings in Bioinformatics 22:bbab102.

Karcher MD, Palacios JA, Bedford T, Suchard MA, Minin VN. 2016. Quantifying and Mitigating the Effect of Preferential Sampling on Phylodynamic Inference.Kosakovsky Pond SL, editor. PLoS Comput Biol 12:e1004789.

Katoh K. 2002. MAFFT: a novel method for rapid multiple sequence alignment based on fast Fourier transform. Nucleic Acids Research 30:3059–3066.

Katoh K, Standley DM. 2013. MAFFT Multiple Sequence Alignment Software Version 7: Improvements in Performance and Usability. Molecular Biology and Evolution 30:772–780.

Khare S, Gurry C, Freitas L B Schultz M, Bach G, Diallo A, Akite N, Ho J, Tc Lee R, Yeo W, et al. 2021. GISAID’s Role in Pandemic Response. China CDC Weekly 3:1049–1051.

Kluyver Thomas, Ragan-Kelley Benjamin, Pérez Fernando, Granger Brian, Bussonnier Matthias, Frederic Jonathan, Kelley Kyle, Hamrick Jessica, Grout Jason, Corlay Sylvain, et al. 2016. Jupyter Notebooks—a publishing format for reproducible computational workflows. In: Positioning and Power in Academic Publishing: Players, Agents and Agendas. IOS Press.

Kraemer MUG, Hill V, Ruis C, Dellicour S, Bajaj S, McCrone JT, Baele G, Parag KV, Battle AL, Gutierrez B, et al. 2021. Spatiotemporal invasion dynamics of SARS-CoV-2 lineage B.1.1.7 emergence. Science 373:889–895.

Krambrich J, Mihalič F, Gaunt MW, Bohlin J, Hesson JC, Lundkvist Å, De Lamballerie X, Li C, Shi W, Pettersson JH-O. 2024. The evolutionary and molecular history of a chikungunya virus outbreak lineage.Makalowski W, editor. PLoS Negl Trop Dis 18:e0012349.

Larsson A. 2014. AliView: a fast and lightweight alignment viewer and editor for large datasets. Bioinformatics 30:3276–3278.

Lemey P, Ruktanonchai N, Hong SL, Colizza V, Poletto C, Van Den Broeck F, Gill MS, Ji X, Levasseur A, Oude Munnink BB, et al. 2021. Untangling introductions and persistence in COVID-19 resurgence in Europe. Nature 595:713–717.

Ligges U, Mächler M. 2003. scatterplot3d -An R Package for Visualizing Multivariate Data. J. Stat. Soft. [Internet] 8. Available from: http://www.jstatsoft.org/v08/i11/

Lozano-Fernandez J. 2022. A Practical Guide to Design and Assess a Phylogenomic Study. Pisani D, editor. Genome Biology and Evolution 14:evac129.

McKinney W. 2010. Data Structures for Statistical Computing in Python. In: Austin, Texas. p. 56–61. Available from: https://doi.curvenote.com/10.25080/Majora-92bf1922-00a

Naveca FG, Almeida TAPD, Souza V, Nascimento V, Silva D, Nascimento F, Mejía M, Oliveira YSD, Rocha L, Xavier N, et al. 2024. Human outbreaks of a novel reassortant Oropouche virus in the Brazilian Amazon region. Nat Med 30:3509–3521.

Nunes MRT, Faria NR, De Vasconcelos JM, Golding N, Kraemer MU, De Oliveira LF, Azevedo RDSDS, Da Silva DEA, Da Silva EVP, Da Silva SP, et al. 2015. Emergence and potential for spread of Chikungunya virus in Brazil. BMC Med 13:102.

Paradis E, Schliep K. 2019. ape 5.0: an environment for modern phylogenetics and evolutionary analyses in R.Schwartz R, editor. Bioinformatics 35:526–528.

Pekar JE, Lytras S, Ghafari M, Magee AF, Parker E, Wang Y, Ji X, Havens JL, Katzourakis A, Vasylyeva TI, et al. 2025. The recency and geographical origins of the bat viruses ancestral to SARS-CoV and SARS-CoV-2. Cell 188:3167-3183.e18.

Posit Team. 2026. RStudio: Integrated Development Environment for R. Available from: http://www.posit.co

Price MN, Dehal PS, Arkin AP. 2010. FastTree 2 – Approximately Maximum-Likelihood Trees for Large Alignments.Poon AFY, editor. PLoS ONE 5:e9490.

Purcell S, Neale B, Todd-Brown K, Thomas L, Ferreira MAR, Bender D, Maller J, Sklar P, De Bakker PIW, Daly MJ, et al. 2007. PLINK: A Tool Set for Whole-Genome Association and Population-Based Linkage Analyses. The American Journal of Human Genetics 81:559–575.

R Core Team. 2025. R: A Language and Environment for Statistical Computing. Available from: https://www.r-project.org

Rambaut A, Lam TT, Max Carvalho L, Pybus OG. 2016. Exploring the temporal structure of heterochronous sequences using TempEst (formerly Path-O-Gen). Virus Evol 2:vew007.

Revell LJ. 2024. phytools 2.0: an updated R ecosystem for phylogenetic comparative methods (and other things). PeerJ 12:e16505.

Rieux A, Balloux F. 2016. Inferences from tip-calibrated phylogenies: a review and a practical guide. Molecular Ecology 25:1911–1924.

Robinson DF, Foulds LR. 1981. Comparison of phylogenetic trees. Mathematical Biosciences 53:131–147.

Rossum G van, Drake FL. 2010. The Python language reference. Release 3.0.1. Hampton, NH: Python Software Foundation

Salfati EL, Spencer EG, Topol SE, Muse ED, Rueda M, Lucas JR, Wagner GN, Campman S, Topol EJ, Torkamani A. 2019. Re-analysis of whole-exome sequencing data uncovers novel diagnostic variants and improves molecular diagnostic yields for sudden death and idiopathic diseases. Genome Med 11:83.

Sanderson T, Barrett JC. 2021. Variation at Spike position 142 in SARS-CoV-2 Delta genomes is a technical artifact caused by dropout of a sequencing amplicon. Wellcome Open Res 6:305.

Santa Ardisson J, Vedovatti Monfardini Sagrillo M, Ramos Athaydes B, Corredor Vargas AM, Torezani R, Ribeiro-Rodrigues R, Cruz Spano L, Gaburro Paneto G, Delatorre E, Ventorin Von Zeidler S, et al. 2025. Comparative spatial–temporal analysis of SARS-CoV-2 lineages B.1.1.33 and BQ.1.1 Omicron variant across pandemic phases. Sci Rep 15:10319.

Schliep K, Potts AJ, Morrison DA, Grimm GW. 2017. Intertwining phylogenetic trees and networks. Fitzjohn R, editor. Methods Ecol Evol 8:1212–1220.

Schliep KP. 2011. phangorn: Phylogenetic analysis in R. Bioinformatics 27:592–593.

Schriml LM, Chuvochina M, Davies N, Eloe-Fadrosh EA, Finn RD, Hugenholtz P, Hunter CI, Hurwitz BL, Kyrpides NC, Meyer F, et al. 2020. COVID-19 pandemic reveals the peril of ignoring metadata standards. Sci Data 7:188.

Slowikowski K. 2016. ggrepel: Automatically Position Non-Overlapping Text Labels with “ggplot2.” :0.9.8. Available from: https://CRAN.R-project.org/package=ggrepel

Souza TML, Vieira YR, Delatorre E, Barbosa-Lima G, Luiz RLF, Vizzoni A, Jain K, Miranda MM, Bhuva N, Gogarten JF, et al. 2019. Emergence of the East-Central-South-African genotype of Chikungunya virus in Brazil and the city of Rio de Janeiro may have occurred years before surveillance detection. Sci Rep 9:2760.

Steel MA, Penny D. 1993. Distributions of Tree Comparison Metrics--Some New Results. Systematic Biology 42:126–141.

Turakhia Y, De Maio N, Thornlow B, Gozashti L, Lanfear R, Walker CR, Hinrichs AS, Fernandes JD, Borges R, Slodkowicz G, et al. 2020. Stability of SARS-CoV-2 phylogenies. Barsh GS, editor. PLoS Genet 16:e1009175.

Vakulenko Y, Deviatkin A, Lukashev A. 2019. The Effect of Sample Bias and Experimental Artefacts on the Statistical Phylogenetic Analysis of Picornaviruses. Viruses 11:1032.

Venables WN, Ripley BD. 2002. Modern applied statistics with S. 4th ed. New York: Springer

Villabona-Arenas ChJ, Hanage WP, Tully DC. 2020. Phylogenetic interpretation during outbreaks requires caution. Nat Microbiol 5:876–877.

Vilsker M, Moosa Y, Nooij S, Fonseca V, Ghysens Y, Dumon K, Pauwels R, Alcantara LC, Vanden Eynden E, Vandamme A-M, et al. 2019. Genome Detective: an automated system for virus identification from high-throughput sequencing data. Birol I, editor. Bioinformatics 35:871–873.

Volk SM, Chen R, Tsetsarkin KA, Adams AP, Garcia TI, Sall AA, Nasar F, Schuh AJ, Holmes EC, Higgs S, et al. 2010. Genome-Scale Phylogenetic Analyses of Chikungunya Virus Reveal Independent Emergences of Recent Epidemics and Various Evolutionary Rates. J Virol 84:6497–6504.

Weaver SC. 2014. Arrival of Chikungunya Virus in the New World: Prospects for Spread and Impact on Public Health. PLoS Negl Trop Dis 8:e2921.

Wickham H. 2009. stringr: Simple, Consistent Wrappers for Common String Operations. :1.6.0. Available from: https://CRAN.R-project.org/package=stringr

Wickham H. 2016. ggplot2: elegant graphics for data analysis. Second edition. Cham: Springer international publishing

Wickham H, Bryan J. 2015. readxl: Read Excel Files. :1.4.5. Available from: https://CRAN.R-project.org/package=readxl

Wickham H, François R, Henry L, Müller K, Vaughan D. 2014. dplyr: A Grammar of Data Manipulation. :1.2.1. Available from: https://CRAN.R-project.org/package=dplyr

Wickham H, Hester J, Bryan J. 2015. readr: Read Rectangular Text Data. :2.2.0. Available from: https://CRAN.R-project.org/package=readr

Wickham H, Pedersen TL, Seidel D. 2011. scales: Scale Functions for Visualization. :1.4.0. Available from: https://CRAN.R-project.org/package=scales

Wickham H, Vaughan D, Girlich M. 2014. tidyr: Tidy Messy Data. :1.3.2. Available from: https://CRAN.R-project.org/package=tidyr

Wilke CO. 2015. cowplot: Streamlined Plot Theme and Plot Annotations for “ggplot2.” :1.2.0. Available from: https://CRAN.R-project.org/package=cowplot

Worobey M, Watts TD, McKay RA, Suchard MA, Granade T, Teuwen DE, Koblin BA, Heneine W, Lemey P, Jaffe HW. 2016. 1970s and ‘Patient 0’ HIV-1 genomes illuminate early HIV/AIDS history in North America. Nature 539:98–101.

Wu F, Zhao S, Yu B, Chen Y-M, Wang W, Song Z-G, Hu Y, Tao Z-W, Tian J-H, Pei Y-Y, et al. 2020. A new coronavirus associated with human respiratory disease in China. Nature 579:265–269.

Ye A, Wang L, Zhao L, Ke J, Wang W, Liu Q. 2021. RapidFuzz: Accelerating fuzzing via Generative Adversarial Networks. Neurocomputing 460:195–204.

Zhou P, Yang X-L, Wang X-G, Hu B, Zhang L, Zhang W, Si H-R, Zhu Y, Li B, Huang C-L, et al. 2020. A pneumonia outbreak associated with a new coronavirus of probable bat origin. Nature 579:270–273.

