## Supplementary Material for "Better data, better trees: GenBank-GISAID deduplication and source-specific artifact masking in viral genomics"

To investigate whether the patterns of laboratory-specific bias were consistent across distinct evolutionary branches, we stratified the primary CHIKV-ECSA dataset (n = 8,617) according to the three major sublineages described by Krambrich et al. (2024), namely ECSA-IOL, ECSA-1 and ECSA-2(Krambrich et al. 2024).

We rooted the Baseline tree using the R packages “ape” v5.8.1 (Paradis and Schliep 2019) and “lubridate” v1.9.5 (Grolemund and Wickham 2011). We then generated the corresponding sequence alignment, phylogenetic tree, and metadata table for each monophyletic group, comprising ECSA-IOL (n = 936), ECSA-1 (n = 13) and ECSA-2 (n = 7,668). The sequence alignments were subset from Baseline alignment using SeqKit v1.5 (Shen et al. 2016). The phylogenetic trees were pruned from rooted Baseline tree using the R package “ape” v5.8.1 (Paradis and Schliep 2019).

Each sublineage dataset was processed following the same procedures described for the full CHIKV-ECSA dataset, including LSBFILT, evaluation of 80 masking strategies, root-to-tip regression, selection of the top three candidates, and identification of the optimal masking strategy after outlier removal.

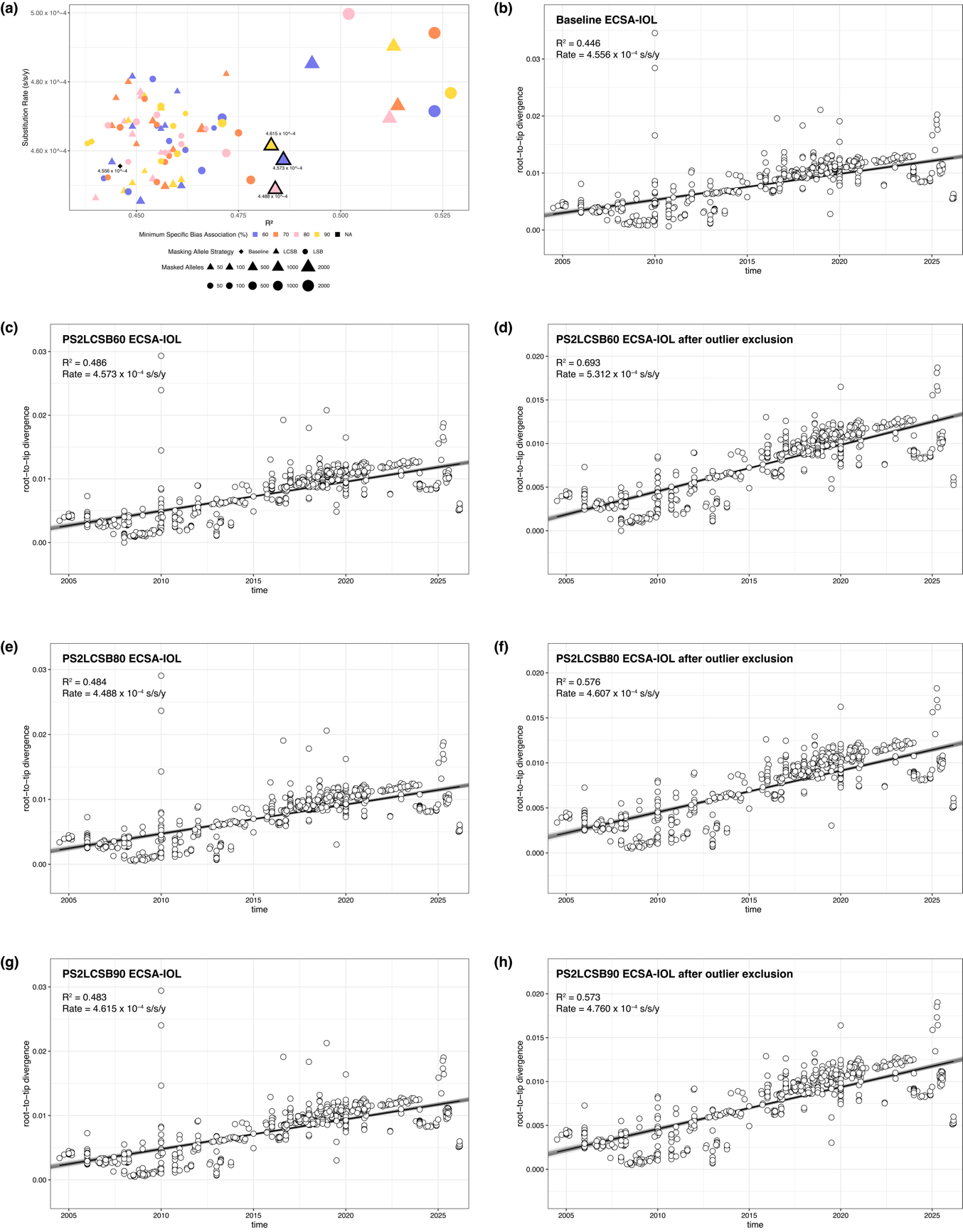

**Figure S1. Effect of allele masking strategy in the ECSA-IOL dataset.** (A) Relationship between root-to-tip temporal signal (coefficient of determination R² from the root-to-tip regression) and substitution rate across 80 masking strategies applied to the ECSA-IOL dataset. Point shape indicates the masking criterion: LSB (laboratory-specific bias, circles), LCSB (laboratory-country-specific bias, triangles), and Baseline (unmasked alignment, diamond). Point colour denotes the minimum specific bias association threshold (blue, 60%; orange, 70%; pink, 80%; yellow, 90%). Point size reflects the number of masked alleles. The top three masking strategies selected for downstream analysis are outlined in black. (B-H) Root-to-tip genetic divergence as a function of sampling time for Baseline (B), PS2LCSB60 (C, before outlier removal; D, after), PS2LCSB80 (E, before; F, after), and PS2LCSB90 (G, before; H, after), with each point representing a sequence, the black line showing the linear regression, and gray lines indicating the confidence interval. Outliers with studentized residuals > 3 were removed prior to regression.

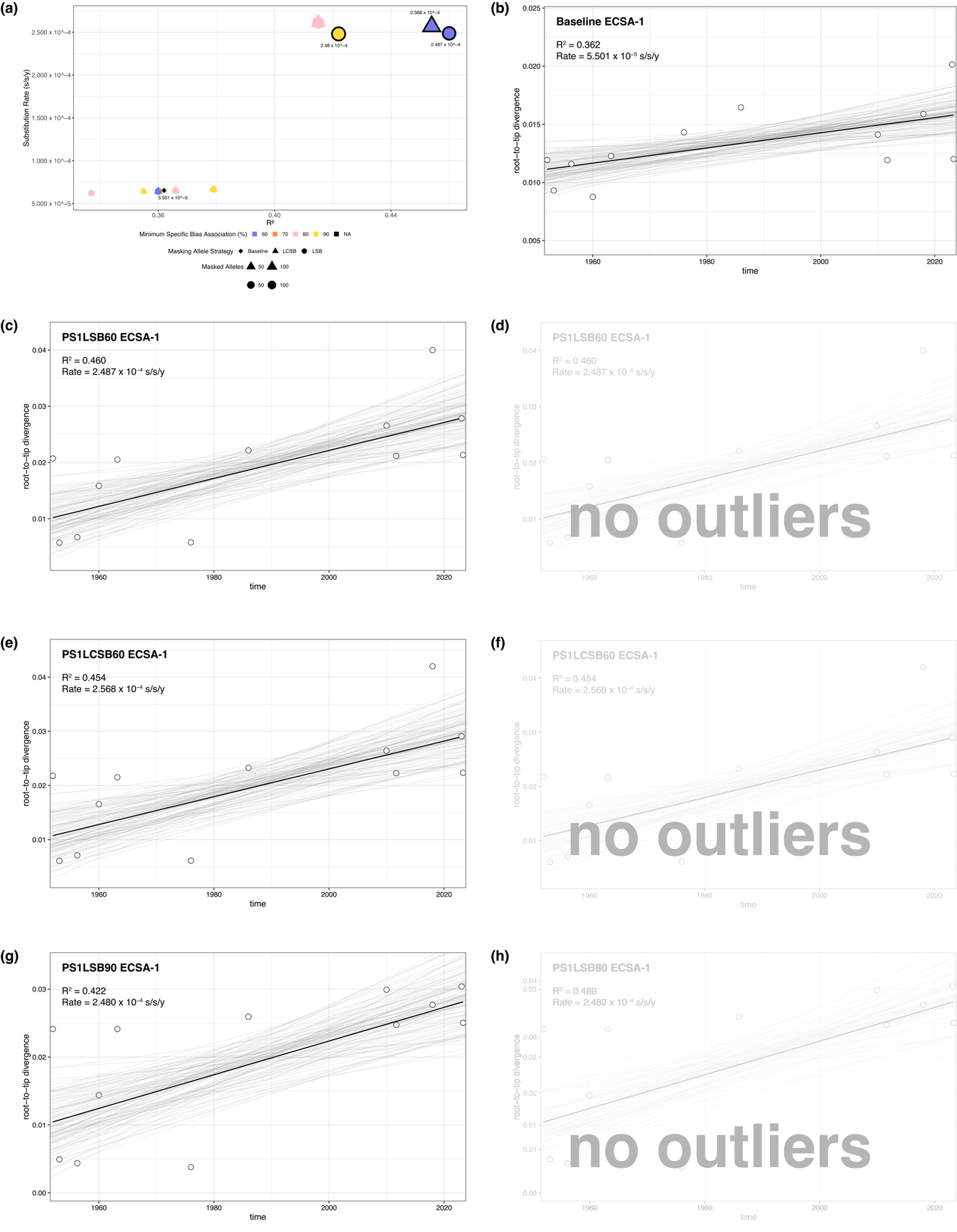

**Figure S2. Effect of allele masking strategy in the ECSA-1 dataset.** (A) Relationship between root-to-tip temporal signal (coefficient of determination R² from the root-to-tip regression) and substitution rate across 80 masking strategies applied to the ECSA-1 dataset. Point shape indicates the masking criterion: LSB (laboratory-specific bias, circles), LCSB (laboratory-country-specific bias, triangles), and Baseline (unmasked alignment, diamond). Point colour denotes the minimum specific bias association threshold (blue, 60%; orange, 70%; pink, 80%; yellow, 90%). Point size reflects the number of masked alleles. The top three masking strategies selected for downstream analysis are outlined in black. (B-H) Root-to-tip genetic divergence as a function of sampling time for Baseline (B), PS1LSB60 (C, before outlier removal; D, after), PS1LCSB60 (E, before; F, after), and PS1LSB90 (G, before; H, after), with each point representing a sequence, the black line showing the linear regression, and gray lines indicating the confidence interval. Due to the limited sample size of the ECSA-1 sublineage (n = 13), no sequences met the outlier threshold (studentized residuals > 3) in any masking strategy.

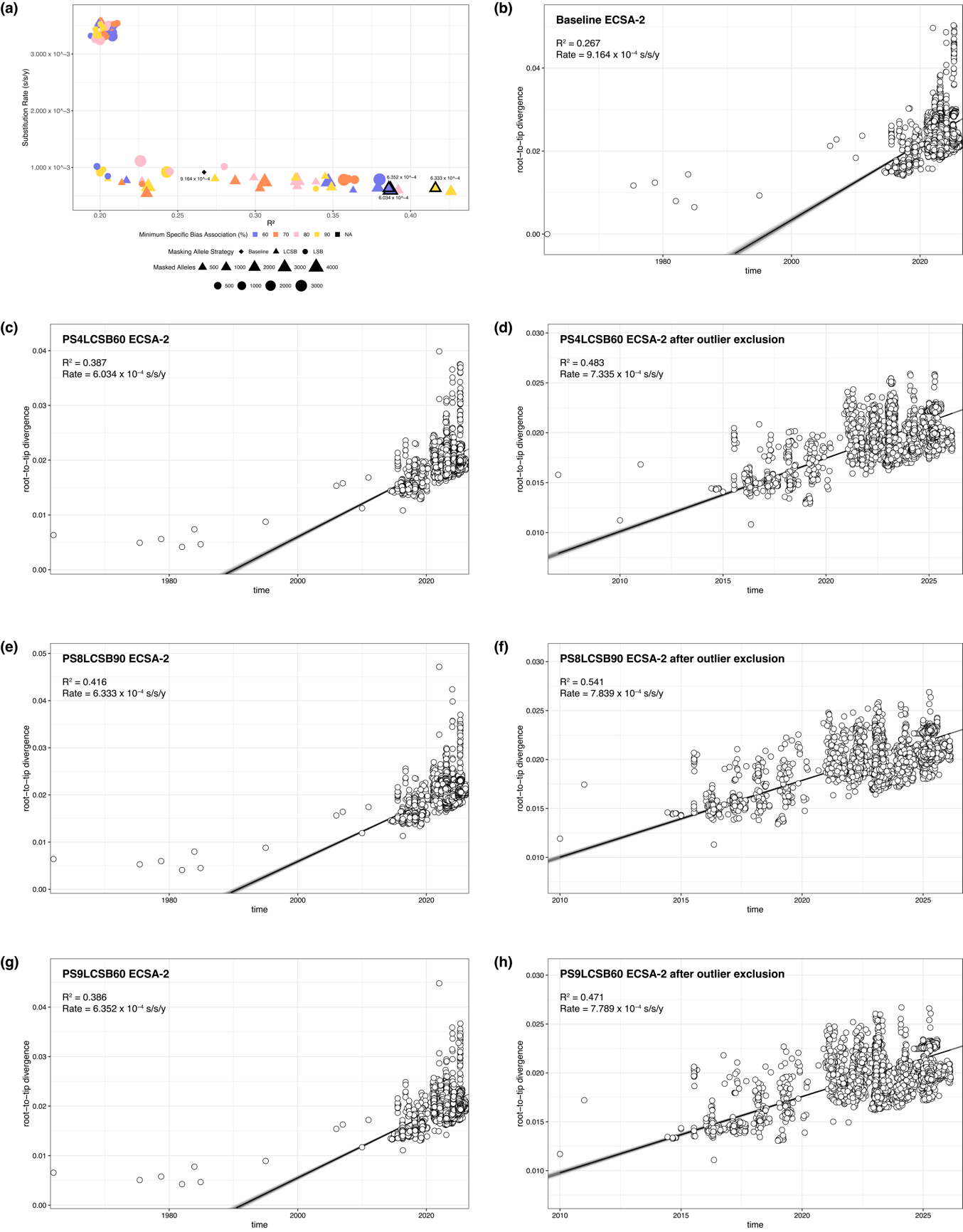

**Figure S3. Effect of allele masking strategy in the ECSA-2 dataset.** (A) Relationship between root-to-tip temporal signal (coefficient of determination R² from the root-to-tip regression) and substitution rate across 80 masking strategies applied to the ECSA-2 dataset. Point shape indicates the masking criterion: LSB (laboratory-specific bias, circles), LCSB (laboratory-country-specific bias, triangles), and Baseline (unmasked alignment, diamond). Point colour denotes the minimum specific bias association threshold (blue, 60%; orange, 70%; pink, 80%; yellow, 90%). Point size reflects the number of masked alleles. The top three masking strategies selected for downstream analysis are outlined in black. (B-H) Root-to-tip genetic divergence as a function of sampling time for Baseline (B), PS4LCSB60 (C, before outlier removal; D, after), PS8LCSB80 (E, before; F, after), and PS9LCSB60 (G, before; H, after), with each point representing a sequence, the black line showing the linear regression, and gray lines indicating the confidence interval. Outliers with studentized residuals > 3 were removed prior to regression.

### References

Grolemund G, Wickham H. 2011. Dates and Times Made Easy with lubridate. *J. Stat. Soft.* 40.

Krambrich J, Mihalič F, Gaunt MW, Bohlin J, Hesson JC, Lundkvist Å, De Lamballerie X, Li C, Shi W, Pettersson JH-O. 2024. The evolutionary and molecular history of a chikungunya virus outbreak lineage.Makalowski W, editor. *PLoS Negl Trop Dis* 18:e0012349.

Paradis E, Schliep K. 2019. ape 5.0: an environment for modern phylogenetics and evolutionary analyses in R.Schwartz R, editor. *Bioinformatics* 35:526–528.

Shen W, Le S, Li Y, Hu F. 2016. SeqKit: A Cross-Platform and Ultrafast Toolkit for FASTA/Q File Manipulation.Zou Q, editor. *PLoS ONE* 11:e0163962.
